# Heritability of complex traits in sub-populations experiencing bottlenecks and growth

**DOI:** 10.1101/2023.11.23.568467

**Authors:** Cameron S. Taylor, Daniel J. Lawson

## Abstract

Populations that have experienced a bottleneck are regularly used in Genome Wide Association Studies (GWAS) to investigate variants associated with complex traits. It is generally understood that these isolated sub-populations may experience high frequency of otherwise rare variants with large effect size, and therefore provide a unique opportunity to study said trait. However, the demographic history of the population under investigation affects all SNPs that determine the complex trait genome-wide, changing its heritability and genetic architecture. We use a simulation based approach to identify the impact of the demographic processes of drift, expansion, and migration on the heritability of complex trait. We show that demography has considerable impact on complex traits. We then investigate the power to resolve heritability of complex traits in GWAS studies subjected to demographic effects. We find that demography is an important component for interpreting inference of complex traits and has a nuanced impact on the power of GWAS. We conclude that demographic histories need to be explicitly modelled to properly quantify the history of selection on a complex trait.

## 2 Introduction

Recent advancements in Genome Wide Association Studies (GWAS) of large human genetic datasets facilitate the study of how genetic variation associates with phenotypes (Hivert et al., 2021, Wang and Huang, 2023, Canela-Xandri et al., 2018), offering insight into the causal genetic structure of complex traits. However, not all populations used in association studies are equivalent - some will contain a diversity of ancestries in their history whilst others may have been relatively isolated. Here we investigate the consequences of studying isolated subpopulations who experienced a population bottleneck on the association with complex traits. Examples include the Ashkenazi Jewish population (Carmi et al., 2014), Icelanders (Rose, 2001), Finns (Kurki et al., 2014), and even experimentally explored in Bacteria (Wein and Dagan, 2019), whose relatively low genetic variation provides a unique opportunity to isolate SNPs associated with traits and disease (Hatzikotoulas et al., 2014). The same principle applies to the ‘Out-of-Africa’ bottleneck that affected all non-Africans, in which includes the largest studies such as UK Biobank (Bycroft et al., 2018) on which — arguably too much (Sirugo et al., 2019) genetic inference is based. However, the genealogies of the ancestral populations in which complex traits evolved may not be well represented by the more homogeneous subgroup that was studied. Here, we explore the consequences of studying these population subgroups via simulation.

A number of Single Nucleotide Polymorphisms (SNPs) contribute to genetic variation of a single complex trait, with a distribution of the effect size *β*_*i*_ for each SNP *i* (Zeng et al., 2018). The frequency of the SNPs *f*_*i*_ can depend on the effect size, which (along with polygenicity and linkage disequilibrium between them) can jointly be considered as the ‘genetic architecture’ of the trait (Johnson et al., 2021). This relationship is linked to whether the trait is under selection, and can be quantified by a selection parameter *s* that takes values *s <* 0 if variation in the trait is selected against (‘negative selection’ which reduces the frequency of large effect SNPs) and *s >* 0 if the trait is selected to change (‘positive selection’ which makes large effect SNPs more common). The complex traits we consider here evolved a long time ago and are now under negative selection. However, the frequency of SNPs is affected by genetic drift, an evolutionary mechanism that leads to changes in the frequency of different alleles between generations (Star and Spencer, 2013), and hence different apparent selection. The relationship between frequency and effect size can be amplified or reduced by drift depending on the genetic architecture of a trait, the history of the population (Ashraf and Lawson, 2021), and which SNP frequencies are examined. Figure 1 illustrates this process.

**Figure 1:**
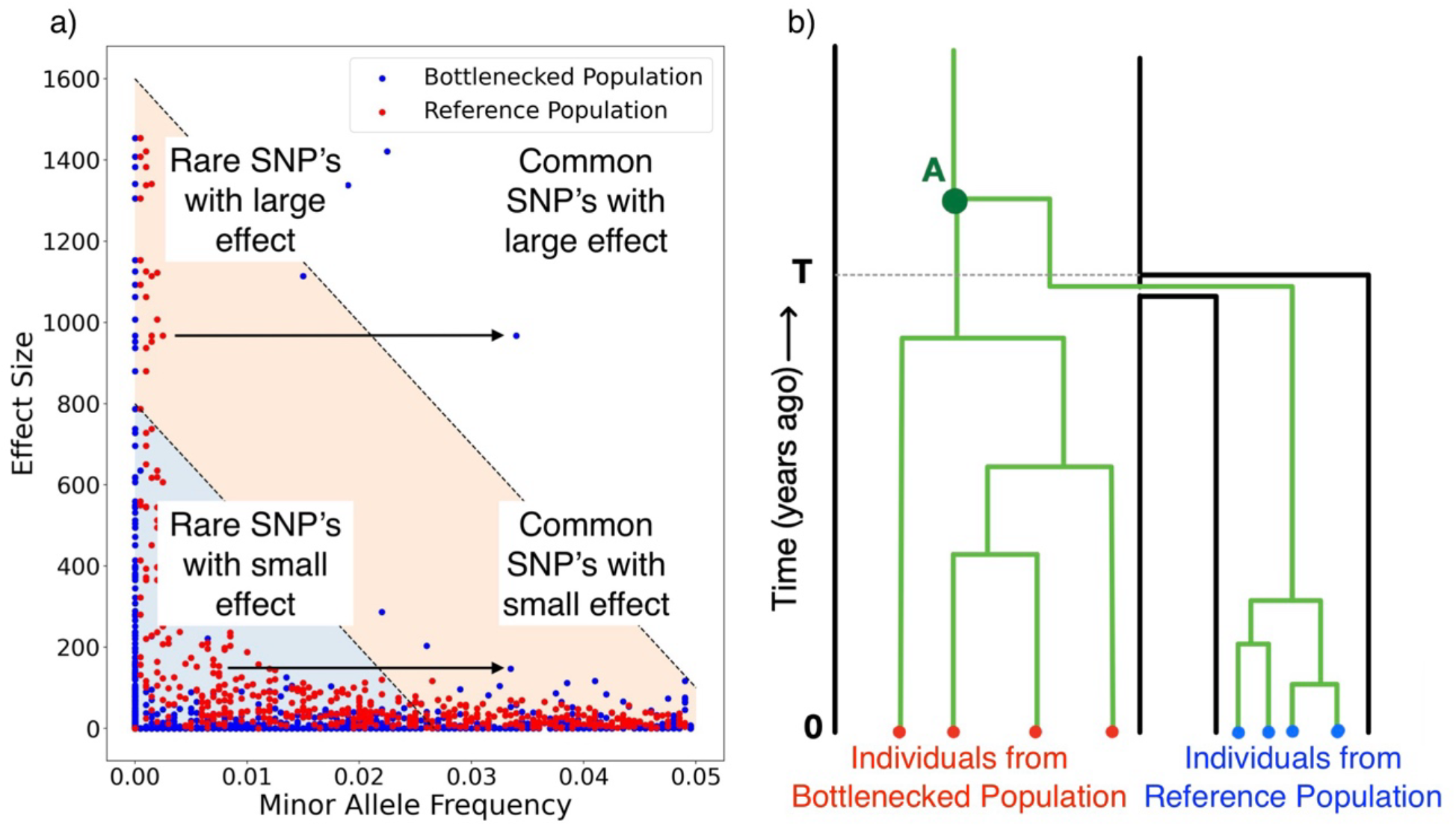
Illustration of the effect of genetic drift on complex trait architecture. a) The relationship between minor allele frequency and the effect size of SNPs in a population that has experienced a bottleneck (blue) and one that has not (red). The arrow shows the effect of drift, allowing rare causal SNPs with large effect size to become common, and changing heritability of complex traits after a population experiences a bottleneck. b) A simple simulation scenario including a population separation time *T* generations in the past, leading to a Reference and Bottlenecked Population of size *N*_0_ and *N* respectively. The green lines show the lineages of individuals for one locus that coalesce at the most recent common ancestor, which are relatively more diverse in the larger population.

Although increased genetic drift is well exploited to discover associations between SNPs and traits, the impact on the distribution of effects is less studied. We consider the impact of a population contraction creating a “bottlenecked population” (potentially considering recovery) within a single ethnicity on the nature of a complex trait. One example is BRCA in Ashkenazi Jewish woman (Metcalfe et al., 2010) for which testing focuses on a single gene, although many genome-wide genetic effects are associated with breast cancer in the wider population (Couch et al., 2013). On a much larger scale, the same bottleneck process leads to benefits and limitations of European Biobanks. The bottleneck that took place as the ancestors of Eurasians left Africa means that these data contain a subset of African diversity, with a concentrated gene pool (Lipson and Reich, 2017). As a consequence, Biobanks of primarily non-African ancestry are not representative of the populations where the traits evolved. Genome-wide trait associations differ systematically in these populations, by the same isolation process described above, though varying in date, severity and migration history. We will investigate how these historical processes affect the association between genetic variation and complex traits.

A natural measure of the genome-wide relationship between genetic variation and a complex trait is heritability *h*_2_, that is, the proportion of phenotype variation due to genetic variation (Wray and Visscher, 2008). Aside from environmental variation, heritability of complex traits is determined by the number and frequency of SNPs and their respective effect sizes. The importance of the increase in frequency of rare SNPs with large effect size due to random drift is demonstrated in Figure 1a in which a population reduction induces a change in the genetic architecture and a subsequent increase in heritability (from 0.5 to around 0.8). Complex traits are inherently polygenic so drift and selection acting on rare SNPs can be offset by that of common SNPs, and vice versa. Therefore, intuition on how genetic drift interacts with selection from well-studied single SNP models, based on the Nearly Neutral Theory of Molecular Evolution (Ohta, 1992) which has been followed up experimentally (Lynch et al., 2016) still require work to translate into the polygenic case. Polygenic selection (Barghi et al., 2020) has been shown to change the distribution of private variants in populations (Durvasula and Lohmueller, 2021) and has been shown to be associated with heritability across traits (O’Connor et al., 2019), (Zeng et al., 2018) but without an explicit accounting for genetic drift.

We use a genome-wide simulation approach to explore how demographic structure, population separation and other factors characterise how a complex trait that has arisen in the past will evolve in future generations. We also consider how the realities of a detection threshold for associations interact with genetic drift. In only well-powered SNP subsets, we find that heritability is still strongly affected by genetic drift, implying that the patterns we observe will impact complex trait analysis from GWAS.

## 3 Results

Our evolutionary framework regards selection on a complex trait considered to have been under selection in the past. Whilst this selection is strong, it is distributed over many loci leading to “nearly neutral” behaviour for each locus under a range of conditions (e.g. Eyre-Walker 2010). This facilitates a neutral model by assuming that selection has maintained SNPs at or below the threshold where selection would efficiently reduce their frequency (Ohta, 1992). This model is practical for inference and widely used for inference of heritability (e.g. Yang et al., 2012), and is appropriate following a contraction if selection is either negligible (because it is less effective in a smaller or growing population) or absent (because the conditions in modern populations have changed).

We use a simulation-based approach to generate population histories accounting for genetic drift, mutation and recombination using the coalescent simulator msprime (Kelleher and Lohse, 2020). This generates (correlated) genealogies for every genetic loci for each individual of both a reference population and a Bottlenecked population. The population structure is simulated such that the ancestral population splits at some time *T* in the past, taken to be 200 generations in the base case, when some members form a sub-population that remains isolated from the Reference population, as shown in figure 1b. Typically, the size of the Reference population, *N*_0_, is large relative to the Bottlenecked population, *N*, taken to be 10000 and 1000 respectively (unless otherwise stated).

To generate a phenotype, we choose a random sample of SNPs to be causal, and assume that the frequency of these SNPs has been under selection as a result of their effect on the phenotype. A proportion 1 − *π* of SNPs are assumed to have no causal effect whilst the proportion *π* (typically small) that contribute to phenotypic variance are distributed as:

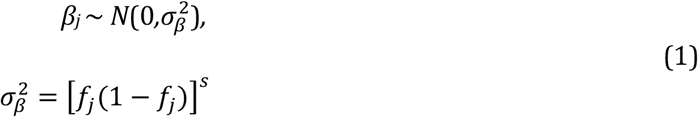

where *f*_*j*_ is the frequency of the *j*th mutation in the sample. Because we are dealing with a simulation in which effect sizes are known, we can use a scaled mutation rate *µ* = *πµ*_0_ as the mutation rate for causal SNPs leading to a random number of causal SNPs *L* with corresponding genotype values *x*_*ij*_ for individual *i* at SNP *j*, where frequencies *f*_*j*_ are for the reference population.

This is used to construct a simulated phenotype 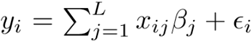 (Zhang et al., 2010). The relationship between effect size and frequency is determined using the parameterisation in Yang et al., (2011) which is equivalent to the model used in LDAK (Speed et al., 2012). Selection strength (on the trait) is parameterised by *s* in Eq. 1, and we report results for negative selection (*s* = −1) so that SNPs with large effect sizes have been selected to become rare. These effect sizes are transferred from the reference to the bottlenecked population, under the assumption of constant environmental variance. The environmental noise *ϵ*_*i*_ is chosen to control the heritability in the reference population (see Methods) to *h*_2_ = 0.5.

Figure 2a-b) shows the distribution of heritability in simulated traits from this model for Bottlenecked Populations of different sizes (from *N* = 500 to 10000) with T=200. With the parameters described, the population contraction typically leads to an decrease in the heritability of complex traits. Several factors influence the extent to which heritability is impacted. A more severe contraction leads to lower levels of heritability on average. This is a result of amplified genetic drift due to fewer founding members of the sub-population, resulting in many SNPs being lost. Conversely, the variance increases with contraction size, as there is a possibility of rare SNPs with large effect being able to reach high prevalence.

**Figure 2:**
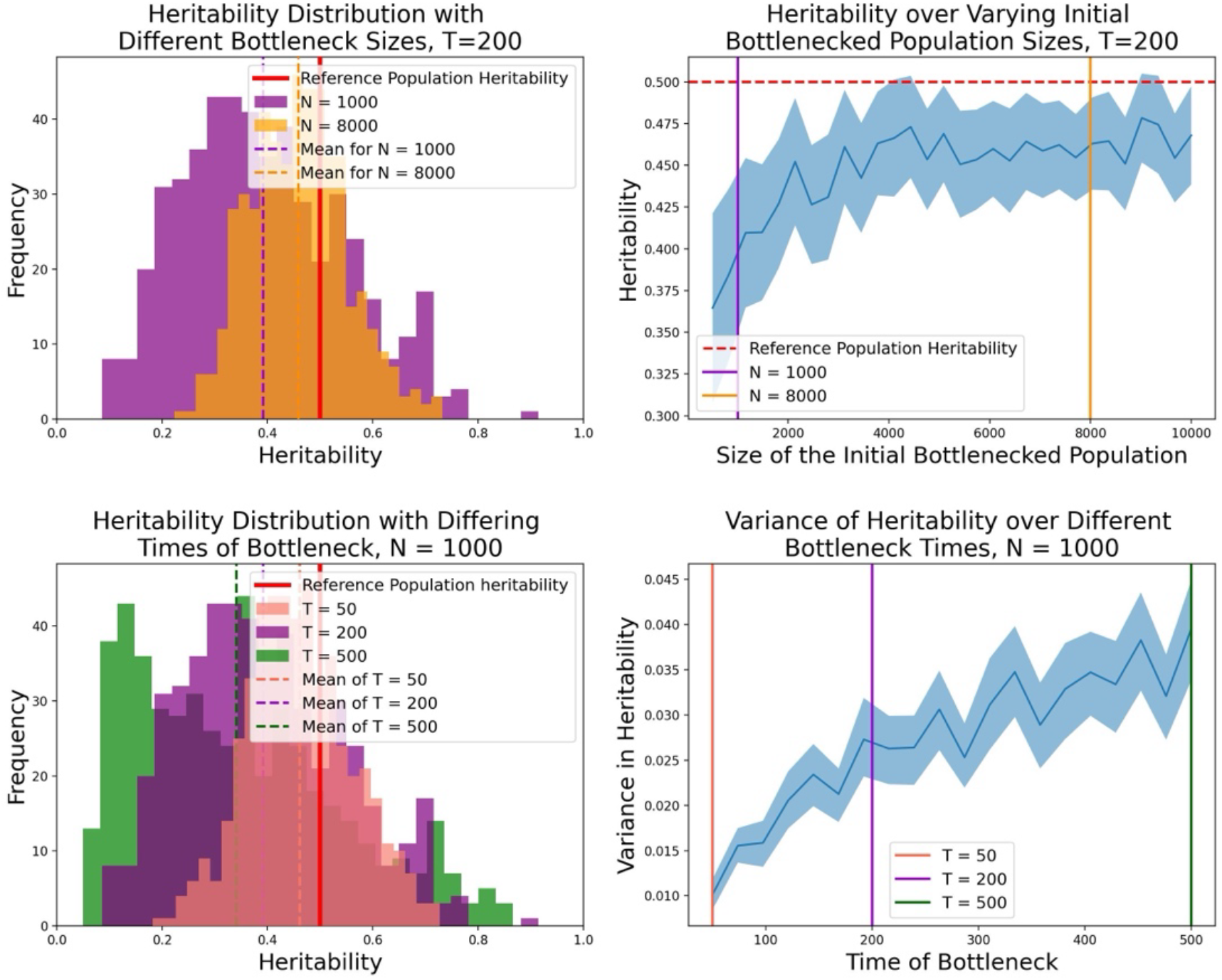
Impact of demographic factors of a bottleneck on heritability. a) Distribution of heritability at different sizes of bottleneck over 500 repeated simulations. Heritability distributions of populations with initial sizes *N* = 1000 and *N* = 8000 with a bottleneck that occurred at *T* = 200 generations ago. b) Mean heritability decreases as the size of the initial Bottlenecked population *N* decreases. 30 samples were taken at each bottleneck size and the mean and error bars are plotted. c) Heritability is distributed with similar tail but different mean and variance over 3 different Bottleneck times. 500 samples are taken for each bottleneck population with different times, *T* = 50,200,500. d) There is a positive relationship between the time in the past that the bottleneck occurred, and the variance in heritability. 95% confidence intervals are shown for the 200 samples taken at each time point using an empirical bootstrap.

The timing of the contraction can also impact how heritability is distributed. The mean heritability of the complex trait in the bottlenecked population decreases with age as more SNPs are likely to be lost. However, traits vary more when the contraction was longer ago (from T=50 to 500, Figure 2c-d) as the most extreme effect size SNPs can become more prevalent as a consequence of genetic drift. Over these time scales, the variance of heritability increases close to linearly with time in the past (Figure 2d), so heritability becomes increasingly varied for complex traits with equal genomic architecture. For reference, 200 generations corresponds to a split time of around 6000 years, similar in scale to the origin of the Ashkenazi Jewish population; 500 generations corresponds to 15kya, around the time of the migration to the Americas; and the out-of-Africa bottleneck took place of the order of 2000 generations (60kya) ago.

Introducing migration rate, *m*, from the reference population to the bottlenecked population offsets the impact of the contraction on the heritability of complex traits (Alcala et al., 2013). There is clear dependence between the time of the contraction and the impact of migration (from m=0-0.1) on mean heritability of traits (Figure 3a), with older events being more effected. Only modest migration (m=0.01 with these parameters) is required to mitigate the effect of the contraction. For some problems even ‘one individual per generation’ (Mills and Allendorf, 1996) is enough migration to offset drift, for complex traits the threshold becomes age-dependent as drift interacts with replenishment of variability.

**Figure 3:**
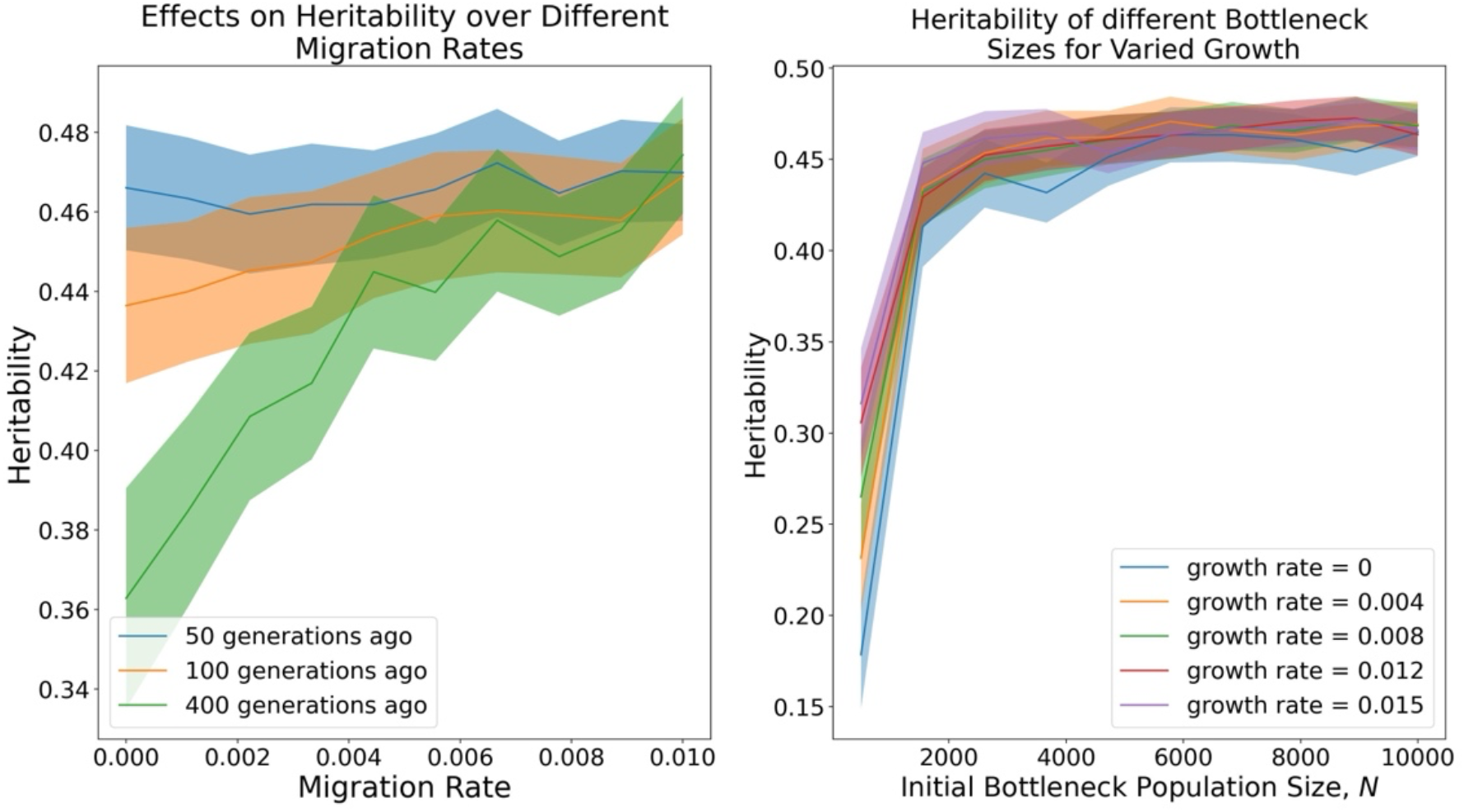
Growth, migration and time since bottleneck change heritability. a) Mean heritability in the Bottlenecked population as migration rate *m* is increased. 400 samples are taken at each time point and the mean with error bars is plotted. b) Effect of bottleneck size and growth on heritability. Shown are the mean and standard errors based on 400 repeated simulations. The reference population is controlled to have heritability 0.5 throughout.

Population growth also impacts heritability change in the population after a contraction. As observed in figure 3b, whilst growth following a bottleneck does recover some heritability, this is dominated by the strength of the contraction. It should be noted that the total bottlenecked population size can considerably exceed the reference population size in these figures - the population grows by a factor of nearly 20 in 200 generations with a growth rate of 0.015.

### 3.1 Impacts for GWAS inference

We have assumed in the simulation study that the effect size of SNPs for a particular complex trait are known. However, in reality effect sizes are inferred from SNP-trait association data using GWAS, which results in constraints being placed on what is known. It might therefore be feared that the outcomes of this paper cannot be directly applied to current complex traits, which we explored in (Ashraf and Lawson, 2021). There are 2 main issues that impact inference in real studies. The first of these is linkage disequilibrium which causes difficulties in isolating causal SNPs. We do not address this problem in this paper but it is explored elsewhere, for example by fine mapping, e.g. in (Malo et al., 2008, Thomas et al., 2011), and is unlikely to prevent the genetic drift we describe here.

The second issue pertains to the lack of statistical power of the data in real GWAS studies, weakening the power to resolve many traits. Whilst the models for which our genetic architecture was introduced (Yang et al., 2011, Speed et al., 2012) already considered missing heritability, these models assume selection acts to create the distribution of effect sizes relating to frequency that are present in the observed population. However, that distribution changes by genetic drift. To explore this further, we considered only SNPs that surpass a threshold of detectability in the bottlenecked population, either by conducting GWAS and ranking by p-value or using variance explained 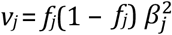 in the trait. We considered only the top SNPs by choosing a variance explained threshold over a large number of simulations. For example, when we report the top 1*/*2_5_ SNPs, we consider the top ∼ 3% of SNPs when averaged over 200 traits. As shown in Figure 4, the heritability distribution can be effected by limited detection power. Specifically, for traits with low heritability in the target population, measured heritability for thresholded SNPs drops, as there are few (or even no) remaining SNPs that have sufficient explanatory power to be recorded. However, for traits with high observed heritability, the true heritability remains high. It follows that our observations on the impact of genetic drift on selection are valid for traits with high heritability, and increasingly for traits with lower heritability as GWAS sample size grows. This also implies that (as is widely assumed) there will be a number of traits that are highly heritable, but that are more difficult to study in drifted (e.g. European) populations.

**Figure 4:**
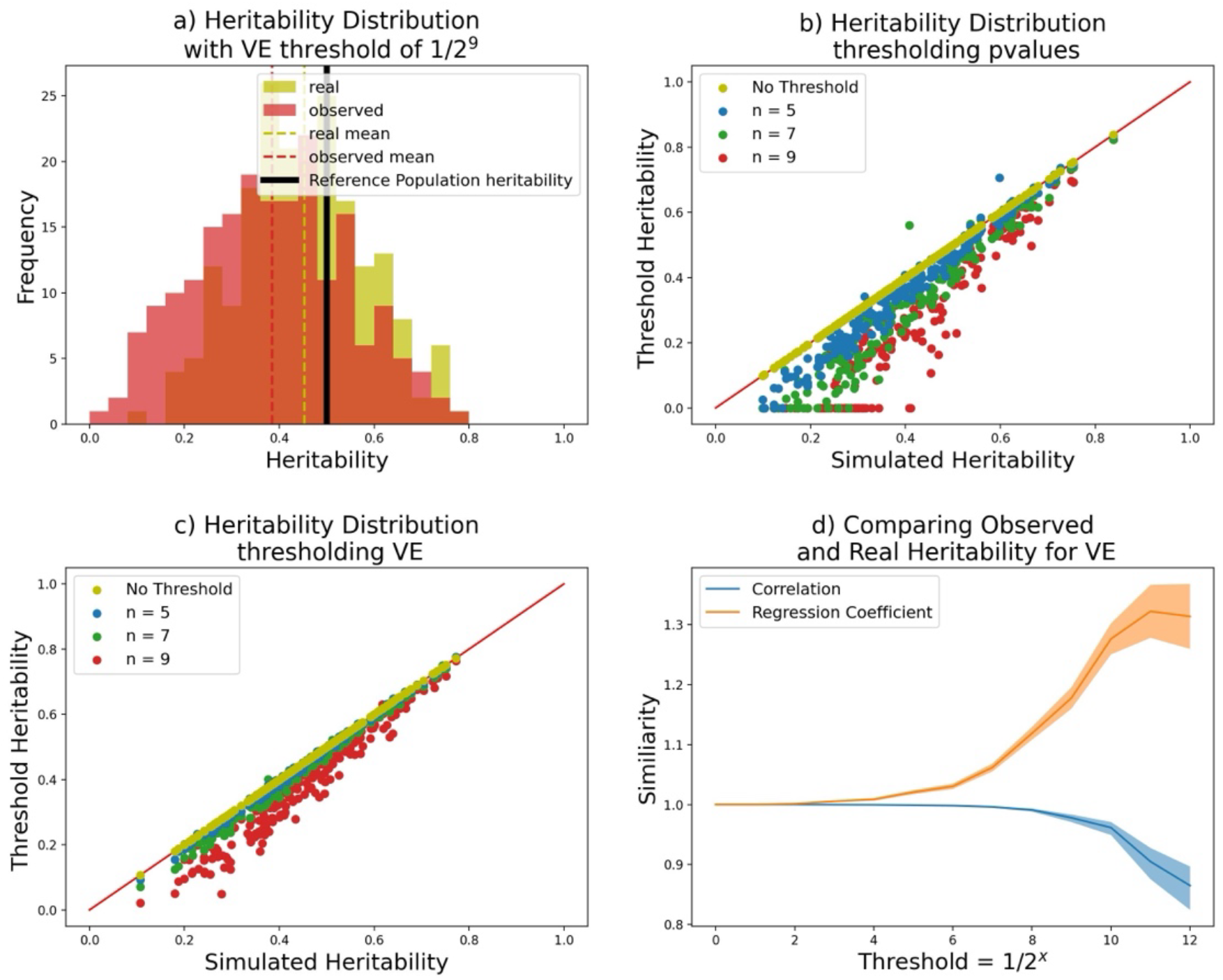
The distribution of heritability of a bottlenecked population as observable in a GWAS study against simulated heritability present, over 200 repeated experiments. a) the simulated heritability distribution (yellow) and observed thresholded heritability (green) where only the 1*/*2^9^ percentile of most informative SNPs (by VE=variance explained) are observed. B) Comparing heritability when thresholding on p-values over 200 repeated experiments, where the threshold heritability only captures the t=1/2^n^ most extreme SNPs. The red line depicts values such that simulated and threshold heritability are equal. c) as b, but where SNPs are ranked by variance explained. D) Pearson correlation and regression coefficient between the real heritability and threshold heritability for thresholds *t* = 1*/*2^*n*^ with standard error bars.

## 4 Discussion

Population bottlenecks lead to genetic drift, which simultaneously reduces genetic variation whilst shifting the frequency of causal variants. Because large effect variants were selected to be rare, genetic drift will tend to increase the variance of these and hence can dramatically increase heritability. However, prediction of heritability is also affected by demographic factors, including growth and migration. Out of this, some general trends emerge.

Sharper bottlenecks have more dramatic impacts on the heritability of complex traits, likely due to a stronger ’founder effect’. The average heritability of a complex trait decreases for population contractions that occurred long ago, as the total amount of genetic drift determines mean heritability. However the variability of heritability between equally selected traits increases as the bottleneck age increases. Therefore some traits increase in heritability and some large effect SNPs rise to high frequency. Migration has a relatively simple impact on heritability as a regression to the mean, by mixing the two populations and maintaining all SNPs at intermediate frequency.

The discoverability of SNPs in GWAS does not appear to strongly interact with the relationship between heritability estimated in a bottlenecked population, and the heritability evolved in the reference population, as long as the heritability was not too low. Therefore inferences from GWAS studies are likely to be effected by the genetic drift process as we described here, even when only a small fraction of the effective SNPs are observed.

The simulation models used here imply that whatever genomic architecture evolved in ancestral populations, the complex process of demography will have changed it. We have demonstrated that it is relatively straight forward to perform inference forward in time, but there are currently no tools to perform the inverse inference: given an observed genomic architecture, and putative population history, can we infer how genomic architecture looked in the past? It seems unlikely that it will resemble what we see today. Further, it would seem to be sensitive to demographic parameters that remain difficult to infer (Lohmueller, 2014). Ancient DNA studies (Irving-Pease et al., 2021) provide constraints on what ancient populations might have looked like, but without massive samples we are not likely to obtain high-quality measurements of the frequency and correlation structure of causal variants. Understanding selection for complex traits then is likely to remain a challenge (Kemper et al., 2014, Hallgrimsson et al., 2014). Quantifying these issues better is important for predicting future evolutionary responses, for example to climate (Wieczynski et al., 2021) and the Anthropocene (Zawada et al., 2019).

## 5 Materials and Methods

### 5.1 Generating Genealogies

Tree sequence data for individuals in the chosen demography is generated using msprime (Kelleher and Lohse, 2020). The demographic model chosen here is outlined in Figure 1b, where two populations are derived from an ancestral population that experiences a split at some point in the past.

For a single simulation, the genealogies are created on 0.1 Morgans of genome. This is kept short for computational reasons, but the 200 simulations we average over can equivalently be thought of as generating 20 Morgans, with no change to the conclusions. However, it is important to emphasise that polygenicity and recombination rate are parameters of genomic architecture (See Supplement on scaling genomes). The time at which this split occurred and size of the initial bottleneck population are parameters that are explored throughout this paper, although a standard of *N*_0_ = 10000 initial people in the Reference population, *N* = 1000 in the Bottleneck population and a split that occurred *T* = 200 generations ago is assumed by default.

Genetic variation is added to the model by superimposing mutations under the infinite sites model, with no time restriction as to when mutations may occur. Other complexities such as a population growth rate and continuous per generation migration rate are included using the corresponding msprime parameters (see Software Availability for exact details).

### 5.2 Complex Trait Simulation

A complex trait is generated by statistically imposing the effect sizes of each SNP present in the reference population. Using the frequency of each SNP in the reference population, *f*_*i*_, the effect sizes *β*_*i*_ are calculated by Equation 1. In this paper the mutation rate is scaled to only include causal SNPs, so all SNPs have non-zero effect sizes.

In order to calculate the phenotype of an individual *i*, we need the contribution of environmental noise, *ϵ*_*i*_. *ϵ* _*i*_ is distributed according to the environmental variance, *V*_*e*_, of the trait, *ϵ*_*i*_ ∼ *N*(0,*V*_*e*_). We simulate *ϵ*_*i*_ with variance chosen such that the heritability in the Reference population remains at 0.5. Heritability is calculated as:

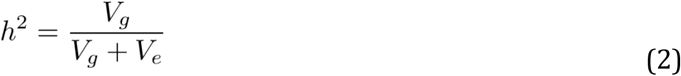

This expression uses the genotype variance of the Reference population, *V*_*g*_ = Var(𝑌^0^), where 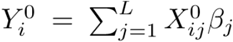 is the genetic component of the phenotype. Here, *X*_*ij*0_ is the genotype of individual *i* at SNP *j* in the Reference population.

Constant environmental variance is assumed in this model, therefore we can formulate the phenotype of for an individual _*i*_ from their genome **X**_*i*_, as well as the *V*_*e*_ and *β* calculated in the reference population. This leads to the mixed linear model

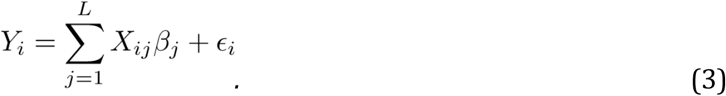

By recalculating the genotype variance *V*_*g*_ in the bottleneck population, the heritability may be calculated using the equation above.

In figures 2-4 new genealogies with a different complex trait (i.e. new effect sizes) were created in each simulation. Figure 2d uses an empirical bootstrap to create error bars for the variance data.

### 5.3 Thresholding

The variance explained by SNP *j* is

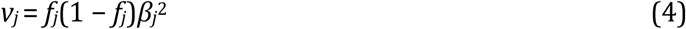

which determines the power to detect the SNP in a GWAS study. We aim to mimic the findings that would realistically be found by thresholding the data to only include SNPs that surpass a certain power threshold.

The power threshold is determined by 200 repeated simulations that share the same demography, but are given unique population genealogies thus different complex trait effect sizes and frequencies. For each simulation the power of each SNP is calculated and the power threshold of the chosen quantile is determined from their combined distribution. In this case the th quantile is used for *n* = 1,2,3,… The complex trait simulation procedure is then repeated, but SNPs that don’t surpass the power threshold in the bottleneck population are discarded. Heritability is then recalculated with the reduced number of effective SNPs.

For Figure 1d, Pearson correlation coefficients between the simulated and thresholded observed heritability values for each threshold level are calculated, as are regression coefficients. Standard error bars are added using Fishers z-transformation.

We further repeated the analysis thresholding not on variance explained, but on p-values, keeping only the smallest.

## Software Availability

The code used to obtain the results in this paper is available at https://github.com/camerontaylor123/TaylorLawson2023.

## Acknowledgements

This work was carried out using the computational facilities of the Advanced Computing Research Centre, University of Bristol - http://www.bristol.ac.uk/acrc/.

## Supplementary Information

The effective amount of genome simulated can be controlled by scaling the simulation parameters, though the scaling is not completely trivial. By changing the recombination rate, sequence length and mutation rate together, we can control the genome wide genomic architecture. For the purpose of this model it is equivalent to changing the polygenicity (i.e. the proportion of SNPs that have an effect on the trait) since in this model all SNPs that are simulated are treated as causal.

Following Kelleher et al., 2016 and Ogundijo and Wang, 2017, the expectation is that if the effective recombination rate, *sequence length* ×*recombination rate* = *constant* and the relative mutation rate, *mutation rate/recombination rate* = *constant*, then the genomic architecture should be equivalent. This follows from the definitions of the scaled recombination rate *ρ* = 4*N*_*e*_*L* and the scaled mutation rate *θ* = 4*N*_*e*_*µ*.

**Figure S1:**
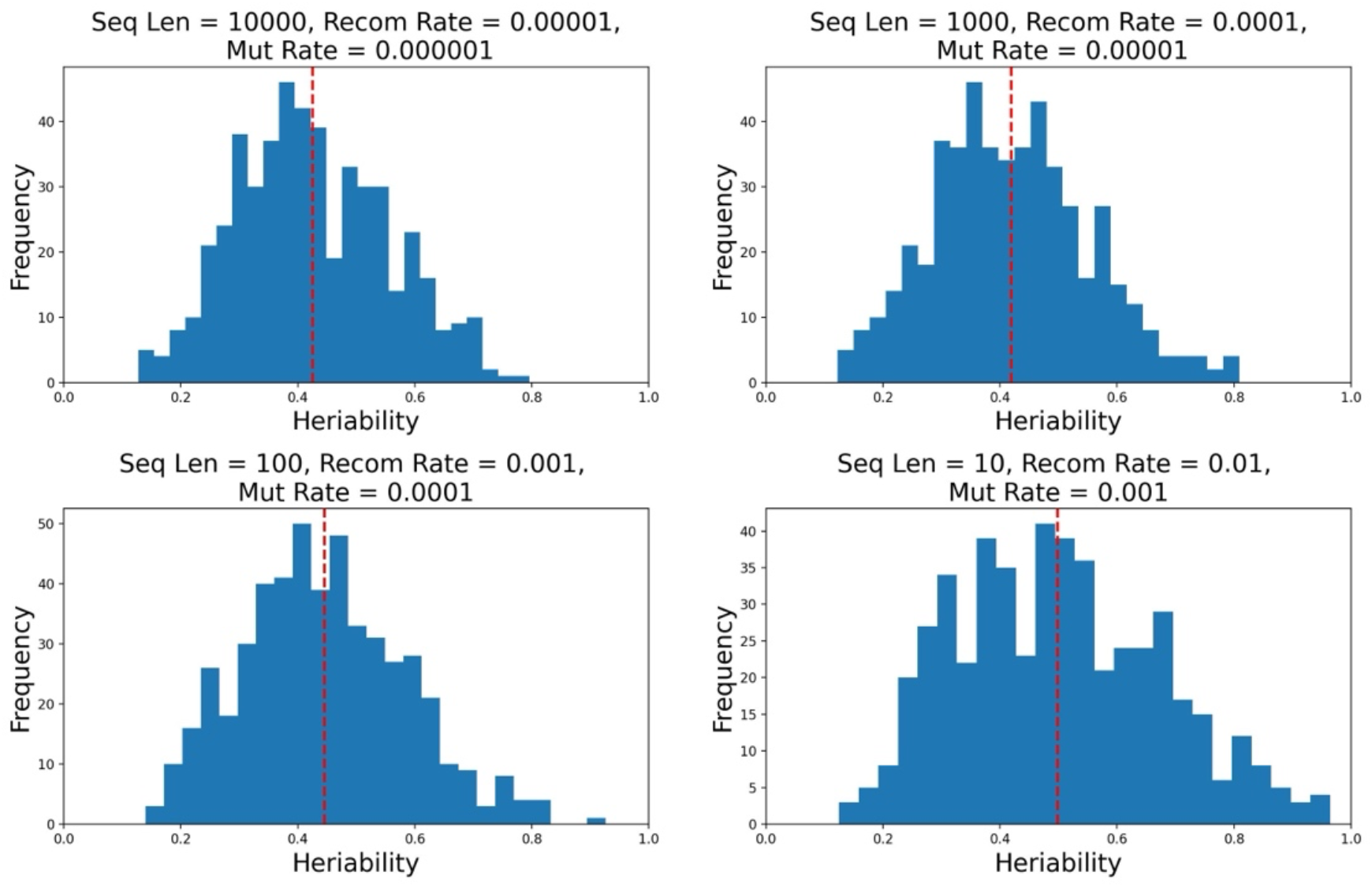
Heritability distribution across scaled sequence length, recombination rate and mutation rate that model the same genetic architecture. (a) sequence length = 1*e*4, recombination rate 1*e*−5, mutation rate = 1*e*−6, (b) sequence length = 1*e*3, recombination rate = 1*e*−4, mutation rate = 1*e* − 5, (c) sequence length = 1*e*2, mutation rate = 1*e* − 3, mutation rate = 1*e* − 4, (d) sequence length = 10, recombination rate = 0.01, mutation rate = 0.001.

Figure S1 shows 500 repeated experiments in four different combinations of heritability and sequence length with *Lρ* = 0.1 and *µ/ρ* = 0.1. The distribution of heritability remains close to consistent in all four cases, though there may be increased volatility as the sequence length is decreased. However, Figure S2 shows the outcome of the experiments with different *recombination rate*, which despite sharing the same expected heritability, produces a different distribution. The variance in heritability increases, as we have decreased the effective number of independent genome regions and hence increased autocorrelation along the genome. In this sense, both polygenicity and recombination rate are parameters of genomic architecture.

**Figure S2:**
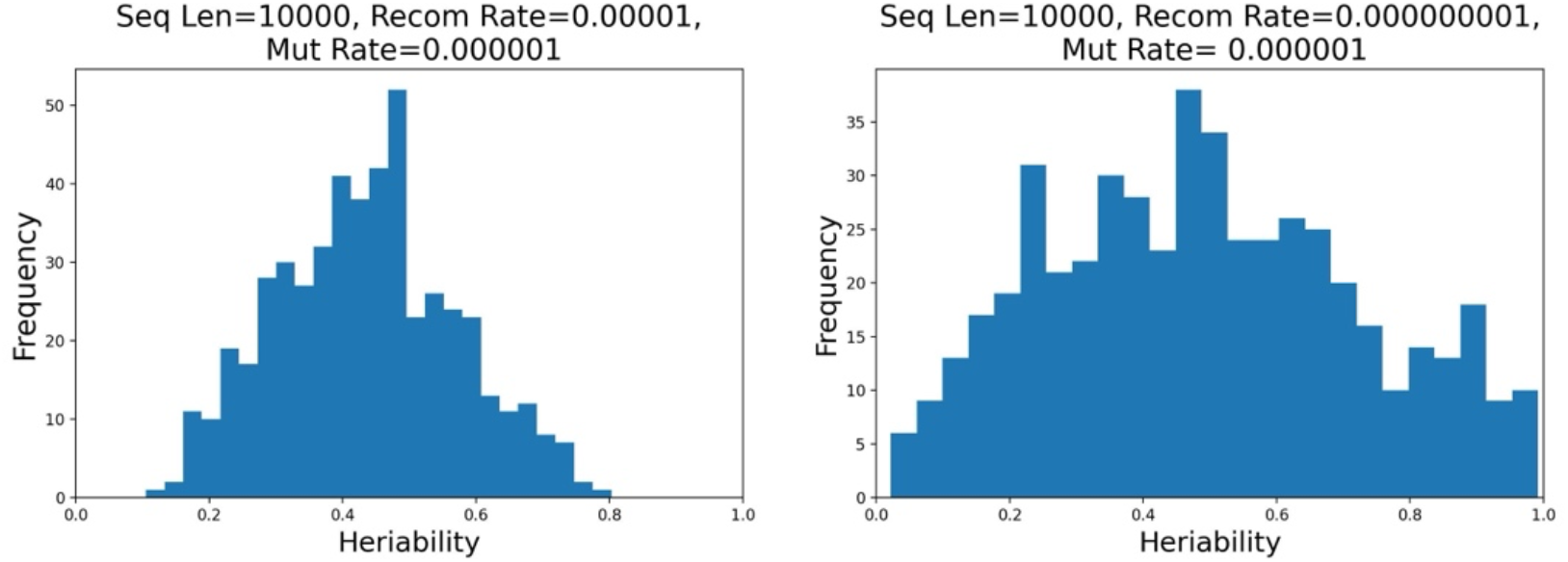
Heritability distribution in differing genetic architectures. (a) sequence length = 1*e*4, recombination rate = 1*e* − 5, mutation rate = 1*e* − 6, (b) sequence length = 1*e*4, recombination rate = 1*e* − 8, mutation rate = 1*e* − 6 (i.e. *ρ* is scaled by 0.001).

